# Quantifying 3D Live-Cell Membrane Dynamics Using Dynamic Metal-Induced Energy Transfer Spectroscopy (dynaMIET)

**DOI:** 10.1101/2024.09.27.614973

**Authors:** José Ignacio Gallea, Narain Karedla, Dongxia Wang, Jörg Enderlein, Tao Chen

## Abstract

The dynamic behavior of cellular membranes underpins essential biological processes, including signal transduction, intracellular trafficking, and mechanotransduction. However, simultaneously quantifying lateral molecular diffusion and vertical membrane fluctuations in live cells remains challenging. Here, we present dynamic metal-induced energy transfer spectroscopy (dynaMIET), which integrates metal-induced energy transfer with fluorescence correlation spectroscopy to resolve three-dimensional membrane dynamics with nanometer axial sensitivity and microsecond temporal resolution. dynaMIET enables concurrent measurement of lateral diffusion and vertical undulations within a single acquisition. We validate the method using simulations and model membranes and demonstrate its robustness in living cells, applying it to the plasma membrane, endoplasmic reticulum, and nuclear envelope. By capturing both molecular mobility and membrane fluctuations, dynaMIET provides a powerful, non-invasive tool for probing membrane mechanics and organization. This advance opens new avenues for studying membrane-associated phenomena in health and disease, including cancer cell mechanics, protein–membrane interactions, and organelle dynamics.

---

Biological membranes are highly dynamic structures composed of a complex mixture of lipids and proteins (*1–3*). Far from being passive barriers, membranes actively regulate a wide range of cellular functions, including adhesion, signal transduction, molecular trafficking, and environmental sensing (*4–7*). These functions critically depend on the dynamic behavior of membrane components. Lipids and proteins constantly move, rotate, and interact within the bilayer, and the membrane itself undergoes spontaneous bending, stretching, and vertical fluctuations (*6, 8*). Abnormalities in these dynamic properties are closely associated with pathological conditions, including cancer (*9, 10*), neurodegenerative diseases (*11, 12*), and immune dysfunction (*13, 14*).

The physical properties of membranes, such as flexibility, tension, and curvature, are closely regulated and contribute to mechanotransduction, vesicle trafficking, and synaptic activity (*6, 15–17*). For instance, the enhanced deformability of cancer cell membranes facilitates metastasis by promoting extravasation through tight endothelial barriers (*9, 18*). In immune cells, membrane mechanics affect the formation of immunological synapses and antigen presentation (*19, 20*). In neurons, changes in membrane composition and fluidity influence vesicle fusion and neurotransmitter release (*6, 21*). Moreover, many viruses, including SARS-CoV-2 and HIV, exploit host membrane dynamics during entry, replication, and egress (*22–24*).

Membrane dynamics span a broad range of spatial and temporal scales. These include molecular rotations of lipids and proteins (*25, 26*), lipid flip-flop between bilayer leaflets (*27, 28*), and lateral diffusion within the membrane plane (*29, 30*), as well as micron-scale vertical undulations and bending (*31, 32*). These dynamic processes are often coordinated with the underlying cytoskeleton (*33*), and their dysregulation can indicate alterations in cell stiffness, metastatic potential, or responsiveness to drugs (*34–36*).

Despite the fundamental importance of membrane dynamics, current experimental tools are limited. Techniques such as fluorescence recovery after photobleaching (*37*), fluorescence correlation spectroscopy (FCS) (*38*), and single-molecule tracking (*39*) effectively measure lateral diffusion of lipids and proteins, but cannot capture vertical fluctuations. Conversely, methods such as flicker spectroscopy (*40*), diffraction phase microscopy (*41, 42*), and reflection interference contrast microscopy (*43–45*) can quantify membrane fluctuations but lack molecular specificity and cannot simultaneously resolve molecular diffusion. Dynamic optical displacement spectroscopy (DODS), an FCS-based method, measures stochastic membrane undulations by positioning the membrane at the confocal focus inflection point, where undulation-induced fluorescence fluctuations dominate while single-molecule diffusion is suppressed through high-density fluorescent labeling (*31*). Recently, employing a similar dense-labeling approach, our group applied metal/graphene-induced energy transfer imaging/spectroscopy (MIET/GIET) (*46–49*) to quantify vertical membrane fluctuations with subnanometer precision (*47*). However, such high labeling densities are difficult to achieve in living cells due to phototoxicity and are incompatible with FCS measurements for diffusion.

Currently, no method exists that can simultaneously capture lateral diffusion and vertical membrane fluctuations both for model membrane and cellular membrane. In this study, we introduce dynamic metal-induced energy transfer spectroscopy (dynaMIET), an approach that simultaneously measure nanoscale molecular diffusion and vertical membrane fluctuations (i.e., three-dimensional (3D) membrane dynamics) with sub-millisecond temporal resolution. The dynaMIET set-up builds on the MIET, an axial super-resolution technique that localizes the molecule with nanometer precision using a thin metal film. By focusing at the metal interface while performing FCS, the detected signal contains contributions from both lateral diffusion of labeled lipids or proteins and vertical membrane undulations, each leaving a distinct signature in the fluorescence intensity autocorrelation. The dynaMIET combines the high axial sensitivity of MIET with the fast, sensitive detection capabilities of FCS to resolve 3D membrane dynamics within a single measurement. This enables accurate quantification at low dye concentrations, compatible with live-cell fluorescent protein imaging, and allows axial membrane fluctuations to be disentangled from lateral molecular diffusion.

To validate our method, we first applied dynaMIET to model giant unilamellar vesicles (GUVs) and benchmarked our results against established techniques. We then extended the method to biologically relevant systems, including the basal plasma membrane (PM), the endoplasmic reticulum (ER), and the nuclear envelope (NE) in both fixed and live cells. These structures play critical roles in processes such as mechanotransduction, protein synthesis, and gene regulation.

## Result

### Principle of dynaMIET for measuring the 3D dynamics of membranes

The 3D membrane dynamics comprises in-plane lateral diffusion and out-of-plane undulations (vertical fluctuations). In microscopic observations, lateral diffusion refers to the in-plane movement of membrane constituents (Fig. 1a), while vertical fluctuations capture out-of-plane undulations and curvature changes of the membrane (Fig. 1a). Our method, dynaMIET, enables simultaneous quantification of both types of dynamics by combining MIET with standard FCS measurements.

**Figure 1:**
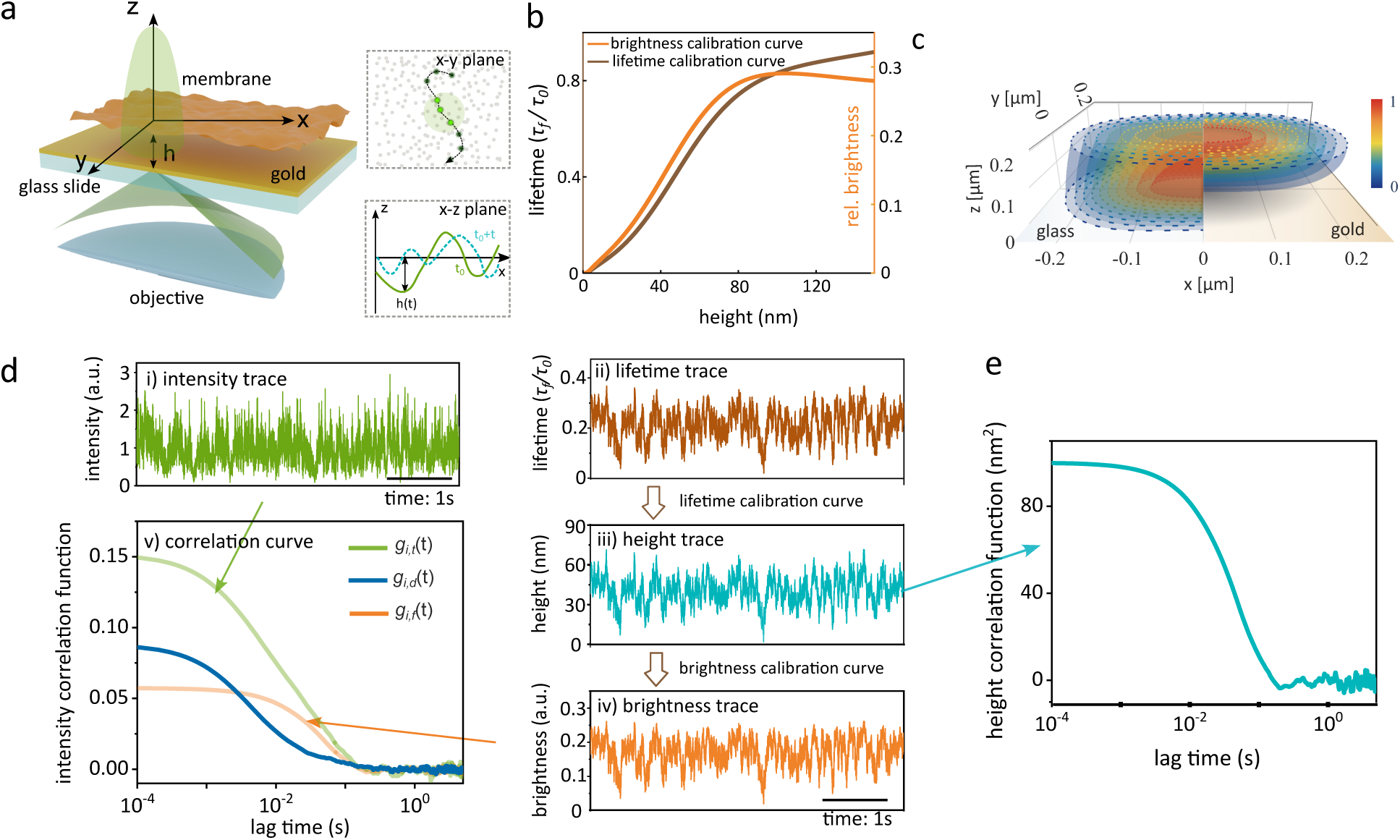
Principle of dynaMIET for measuring membrane dynamics. (a) Sketch of the membrane dynamics within a confocal volume. The coordinate system and notations are defined. The membrane is in the x-y plane and undergoes small displacements in the z direction. Lateral dynamics refer to the diffusion of fluorophores within the x–y plane, while vertical dynamics correspond to the up-and-down fluctuations of the membrane along the z axis. A fluctuating membrane situated above a gold film is schematically depicted. The double-headed arrow indicates the average height *h* of the labelled membrane from the substrate. (b) Calculated relative fluorescence lifetime (*τ*_*f*_ /*τ*_0_) (upper panel) and relative brightness (lower panel) of an emitter as a function of its distance *h* from the surface of a 10 nm gold film. These calculations consider an emitter with a maximum emission wavelength of 680 nm, a fluorescence quantum yield of 0.8, and a random orientation. (c) Comparison of calculated isosurfaces of molecular brightness as a function of position within a confocal focus (N.A. = 1.49) of a linearly polarized beam, with the focal plane positioned at the glass–sample interface, for both a glass substrate and a MIET substrate. (d) Simulation for a fluctuating membrane above a gold surface. Simulated fluorescence intensity trace (i) and lifetime time trace (ii). Calculated height trace (iii) and brightness trace (iv) based on the calibration curves in **b**. (v) Calculated intensity correlation curves. (e) Calculated height correlation curve from **b-iii**.

Fig. 1a illustrates the MIET setup. A typical substrate consists of a glass coverslip coated with a thin gold film. MIET operates via electrodynamic coupling between the excited-state near-field of a fluorophore and surface plasmons in the gold layer. Functionally, the gold film acts as a nonradiative energy acceptor, analogous to the acceptor in Förster resonance energy transfer (FRET), resulting in highly distance-dependent quenching. This coupling modulates both the fluorescence lifetime and emission brightness of the fluorophore as a function of its axial (vertical) distance from the metal surface (see Supplementary Text).

As shown in Fig. 1b, theoretical calculations for a dye whose transition dipole rotates freely and is oriented randomly with respect to the substrate (quantum yield *ϕ* = 0.8, emission wavelength *λ*_em_ = 680 nm) predict that the relative fluorescence lifetime *τ*_*f*_ (*h*)/*τ*_0_ increases monotonically with height (at least over *h* ≲ 150 nm), whereas the brightness *b*(*h*) rises with *h* and reaches a maximum near *h* ≈ 90 nm before decreasing at larger separations. This distance-dependent brightness modulation is critical: within a diffraction-limited excitation focus, lateral diffusion modulates intensity via molecules entering and leaving the excitation focus, whereas the MIET axial dependence makes brightness vary steeply with height over the first few tens of nanometers, as shown in Fig 1c. As a result, small vertical displacements of molecules in a membrane produce large fractional intensity changes, enabling dynaMIET to resolve nanometer-scale vertical membrane fluctuations as well. These axial fluctuations are essentially invisible to conventional FCS, whose axial excitation profile spans ∼ 300 nm and is therefore predominantly sensitive to lateral diffusion within the confocal volume. Therefore, the measured intensity autocorrelation function (iACF) in the presence of a MIET substrate, denoted by *g*_*i*,*t*_ (*t*), reflects both lateral and vertical dynamics:

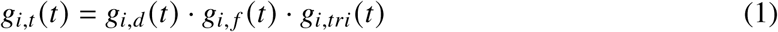

Here, *g*_*i*,*d*_ (*t*) describes lateral diffusion, *g*_*i*, *f*_ (*t*) captures vertical fluctuations, and *g*_*i*,*tri*_ (*t*) accounts for the triplet-state relaxation of the fluorophore. In general, triplet relaxation occurs on the microsecond timescale and does not interfere with the slower membrane dynamics that unfolds on the millisecond timescale. A key challenge here is disentangling the respective contributions of molecular diffusion and vertical membrane fluctuations to the total iACF, which take place on similar timescales. In our previous work, we addressed this by increasing the density of labeling in the bilayers, which suppresses diffusion-related fluctuations due to the larger number of emitters in the detection volume (*47*). However, this strategy inevitably comes at the cost of reducing the access to valuable diffusion information on the constituent proteins or lipids.

To overcome this, we developed a strategy that leverages concurrent determination of the fluorescence lifetime, molecular height, and brightness from the raw photon data. To demonstrate this approach, we simulated the 3D dynamics of a lipid membrane near a MIET substrate as observed in a confocal setup (see Methods for detailed information on the simulations). Figure 1d shows (i) a simulated fluorescence intensity trace used to calculate the total iACF *g*_*i*,*t*_ (*t*) and (ii) a corresponding lifetime trace for a fluctuating membrane. These simulations faithfully replicate the experimental data acquired via time-tagged time-resolved (TTTR) single-photon counting.

We then convert the simulated lifetime trace into a height trajectory *h*(*t*) (Fig.1d-iii) using a calibration curve derived from MIET theory (see Fig.1b). The height trace is further transformed into a corresponding relative brightness trace using the calculated brightness-vs.-height curve (see Fig.1b). This yields a synthetic trace from which we compute the fluctuation-only iACF component *g*_*i*, *f*_ (*t*). Using Eq.(1), we isolate the diffusion-related autocorrelation function *g*_*i*,*d*_ (*t*) by dividing the total iACF by *g*_*i*, *f*_ (*t*) · *g*_*i*,*tri*_ (*t*). The lateral diffusion coefficient *D* is then extracted using established 2D diffusion models applicable to lipid bilayers (*38*).

For a fluctuating membrane, the instantaneous vertical position is time-dependent, and the height trajectory *h*(*t*) can be decomposed into a time-averaged mean height *h*_0_ and the time-dependent instantaneous displacement *δh*(*t*), such that *h*(*t*) = *h*_0_ + *δh*(*t*). The temporal behavior of vertical membrane fluctuations is characterized by the height autocorrelation function (hACF), *g*_*h*_ (*t*) = ⟨*δh*(*t*^′^)*δh*(*t*^′^ + *t*)⟩_*t*_′ (Fig. 1e). The amplitude of *g*_*h*_ (*t*) yields the amplitude of the variation of root mean square fluctuation 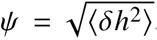, while the characteristic decay time *τ*^∗^ represents the relaxation timescale of vertical fluctuations (*31, 47*). Based on these analyses and fitting, we obtained diffusion coefficient *D* = 4.97 ± 0.15 µm^2^/s (mean ± SD, n = 10 independent simulations) and fluctuation amplitude *ψ* = 9.96 ± 0.03 nm (mean ± SD, n = 10 independent simulations) for the simulated data, which closely match the input values used in the simulation (*D* = 5 µm^2^/s and *ψ* = 10 nm).

When transitioning from simulations to experimental dynaMIET, the data analysis must explicitly account for model experimental photophysics such as photobleaching, triplet-state kinetics, and detector artifacts such as afterpulsing. Additionally, in our work-flow, we couple fluorescence intensity-lifetime information via rigorous fluorescence lifetime fitting. These components are integrated into our custom analysis pipeline, which is described in detail in the Materials and Methods section.

### 3D dynamics of GUV membranes

We validated dynaMIET by measuring membrane undulations and lipid diffusion in the proximal membrane of deflated giant unilamellar vesicles (GUVs), and then compared our measurements with well-established theoretical predictions (*50–53*), previous experimental results (*31,47,50–52*), and a comparison with a planar membrane without bending undulations. GUVs were prepared from 1-stearoyl-2-oleoyl-sn-glycero-3-phosphocholine (SOPC) lipids and fluorescently labeled 1,2- Dipalmitoyl-sn-glycero-3-phosphoethanolamine (DPPE-Atto655) in a labeling ratio of 0.05 mol % by electroformation in a sucrose solution with an osmotic pressure of 230 mOsm/L. For fluorescence measurements, see Fig. 2a, GUVs with an internal sucrose concentration of 230 mOsm/L were immersed in a hypertonic solution of 400 mOsm/L and added above a MIET substrate passivated with bovine serum albumin (BSA). The MIET substrate comprised of a 10 nm thick silica layer and a 10 nm gold film on a glass coverslip. The elevated osmolarity outside the GUVs led to their deflation and reduction of membrane tension, inducing membrane undulations close to the substrate surface (*31, 54*). FCS measurements were performed while the laser was focused at the bottom of the chamber within the contact area of the GUV.

**Figure 2:**
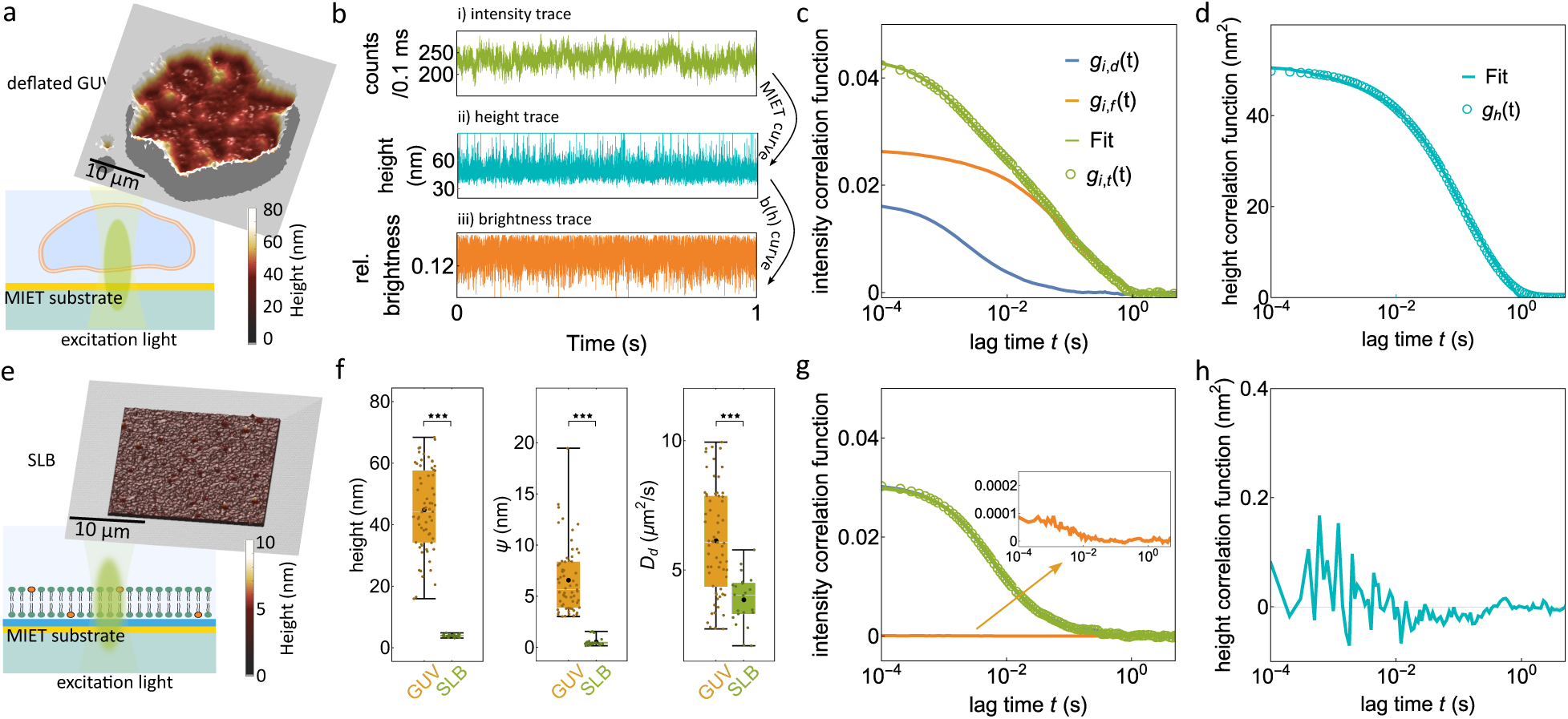
Validation of dynaMIET on model membrane. (a) Schematic of a dynaMIET experiment on a GUV above a MIET substrate. In the upper panel, exemplary 3D reconstructions (height image) of the proximal membrane of a deflated GUV on a MIET substrate are shown, with height encoded by color. (b) Calculated fluorescence intensity time trace (i), height trace (ii), and brightness trace (iii) for the proximal membrane of one deflated GUV. (c) Various fluorescence iACFs derived from measurements of the proximal membrane of surface-immobilized GUVs on a MIET substrate. The green solid line represents a fit with Eq. S3. (d) Calculated hACF (dotted lines) for the same deflated GUV. Fit values obtained from fitting the hACF with eq 2 (solid line) are also shown. (e) Schematic of a dynaMIET experiment on a SLB deposited on a MIET substrate. In the upper panel, exemplary 3D height profiles of the SLB on the MIET substrate. Height values are encoded by color. (f) Box plots for mean height value *h*_0_, height correlation amplitude *ψ*, and diffusion coefficient *D*. Box plots show the 25th–75th quantiles (box), median (white line), mean (black dot), and whiskers (minima to maxima). We performed *n* = 22 independent measurements over 2 independent experiments for SLBs, and *n* = 65 independent measurements over 4 independent experiments for deflated GUVs. Significance levels *P* were evaluated by a Mann-Whitney *U*-test: *P* < 0.001 (★ ★ ★), *P* < 0.01 (★★), *P* < 0.05 (★). (g) Fluorescence iACFs derived from planar SLB supported on a MIET substrate. The green solid line is fitted with eq S3. Inset shows an enlarged plot of the height-fluctuation-related iACF. (h) Calculated hACF (dotted lines) for the same SLB.

GUV deflation was evidenced by the non-circular geometry of the contact area on the MIET substrate (Fig. 2a). Fig. 2b-i, Fig. 2b-ii, and Fig. 2b-iii display the measured intensity trace *I* (*t*), the calculated height trace *h*(*t*), and the brightness trace *br* (*t*), respectively, with a time-bin width of 100 µs, where the mean number of photons per time bin was ∼ 210. Based on the dynaMIET analysis, we successfully separated iACF into contributions from lateral lipid diffusion and from vertical undulations of the membrane. Moreover, we determined the translational diffusion coefficient of membrane lipids as *D* = 6.75 µm^2^/s by fitting the iACF *g*_*i*_ (*t*) with eq. S3. This diffusion coefficient aligns well with values measured for deflated GUVs on a pure glass substrate (Fig. S3), where vertical membrane undulations do not contribute to the fluorescence intensity fluctuations.

Subsequently, we derived an hACF to assess vertical membrane undulations (Fig. 2d). The membrane height fluctuations of the deflated GUVs exhibit a larger amplitude, characterized by a root mean square height fluctuation of *ψ* = 6.6 ± 3.2 nm and a relaxation time of *τ*^∗^ = 59 ± 55 ms (mean ± SD, *n* = 65). This observation is consistent with our previous measurements using a high-label-concentration method (*47*), and aligns well with values obtained by other techniques (*50*).

The measured hACF *g*_*h*_ (*t*) can be effectively modeled using a theoretical framework outlined in Refs. (*50–53*)). The model parameters are the membrane bending rigidity *κ*, the membrane tension *σ*, the effective viscosity *η*, and an interaction potential *γ* that characterizes the coupling between the fluctuating membrane and the substrate. The hACF is modeled by the following equation:

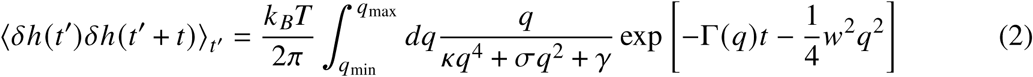

where *w* denotes the diameter of the confocal detection area. The relaxation rate Γ(*q*) includes hydrodynamics and wall effects next to a solid support and is defined by

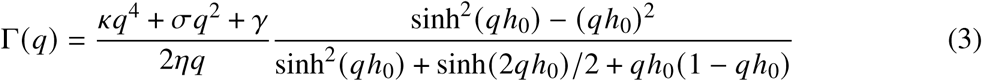

The integration boundaries of the integral in eq. (2) are *q*_min_ = (*γ*/*κ*)^1/4^ and *q*_max_ = 1/*h*_0_ (*53*). When applying this model to describe the hACF of SOPC membranes, we fixed the value of the bending modulus to *κ* = 20 *k*_*B*_*T* as reported in the literature (*31, 55*). We approximated the effective viscosity as the arithmetic mean of the viscosity of buffer solutions inside and outside the GUV, yielding *η* ∼ 1.2 mPas. The diameter of the confocal detection area in our measurement was determined as *w* = 280 nm by measuring the FCS of a sample with a known diffusion coefficient. We acquired 65 hACF measurements and fitted them with the model; a representative hACF curve (same data point for Fig. 2c) and the corresponding fitted curve are shown in Fig. 2d. From these fits, we derived a surface tension value of *σ* = 13 ±12 µJm^-2^ (mean ± SD, n = 65) and an interaction potential strength value of *γ* = 8 ± 6 MJm^-4^ (mean ± SD, n = 65), in excellent agreement with published results for similar GUVs (*47, 50–52*).

As a control, we performed dynaMIET measurements on a supported lipid bilayer (SLB) on a MIET substrate (Fig. 2e). We compared the parameter values obtained from the SLB measurements with those obtained from the deflated GUVs (Fig. 2f). An SLB is a planar membrane on a substrate with only a thin hydration layer of only 1 ∼ 2 nm between the substrate and the membrane (Fig. 2e). Consequently, an SLB can be considered as a membrane without any vertical undulations. As anticipated, the brightness-related iACF is nearly flat, indicating that all intensity fluctuations arise from the lateral diffusion of lipid molecules (Fig. 2g). Similarly, the observed hACF amplitude for an SLB is negligible (*ψ* = 0.54 ± 0.41 nm, mean ± SD, n = 22, Fig. 2h).

The diffusion coefficients for deflated GUVs obtained with dynaMIET and with conventional FCS (measured on a glass substrate for deflated GUVs, see Fig. S3) are very close to each other (dynaMIET: 6.13 ± 2.03 µm^2^/s; conventional FCS: 6.5 ± 2.0 µm^2^/s).The diffusion coefficient *D* = 3.85 ± 0.95 µm^2^/s (*n* = 22) of lipid diffusion in an SLB is considerably slower than that of lipids in a deflated GUV (*D* = 6.13 ± 2.03 µm^2^/s, *n* = 65). This difference is due to the fact that lipid diffusion in the proximal leaflet of an SLB is impeded by a strong interaction between the lipids and the solid support.

The 3D dynamics of the model GUV membrane, as measured with dynaMIET, aligns well with existing theoretical models and provides reliable values for parameters such as diffusion coefficient, undulation amplitudes, membrane tension, and energy dissipation. These results establish dynaMIET as an accurate and robust technique, which we next apply to investigate the more complex membranes of live cells.

### 3D dynamics of the cellular plasma membrane

The plasma membrane (PM) is a very dynamic structure involved in various cellular functions, including cell motility, cell division, or vesicle trafficking. PM undulations span a broad temporal range, with slow actin waves inducing large-wavelength membrane oscillations at cell edges with time durations of seconds. These oscillations involve vertical membrane undulations that range from 100 nm to 10 µ*m* (*56*). In the basal membrane, smaller amplitude undulations are observed, because of the presence of the extracellular matrix (ECM) between the PM and the substrate. Although pronounced membrane undulations are associated with processes such as cell spreading and motility, even subtle, nanometer-scale fluctuations can modulate the spatial organization of molecules within the plasma membrane. These small-amplitude height variations can promote the formation of nanodomains and membrane invaginations, thereby shaping local biochemical organization (*57*).

We applied dynaMIET to resolve the 3D dynamics of the basal membrane COS7 cells cultured on a MIET substrate with a 10 nm silica spacer, where the PM was labeled with the commercial PM stain CellMask Deep Red. dynaMIET measurements were performed on the basal membrane of both living and fixed cells (Fig. 3a). Fig. 3b displays reconstructed 3D height maps of the basal membrane of COS7 cells. As can be seen, the distance between the basal membrane and the substrate varies between 35 nm to 55 nm. We randomly chose different points in different cells for the measurement of the temporal dynamics of PM undulations.

**Figure 3:**
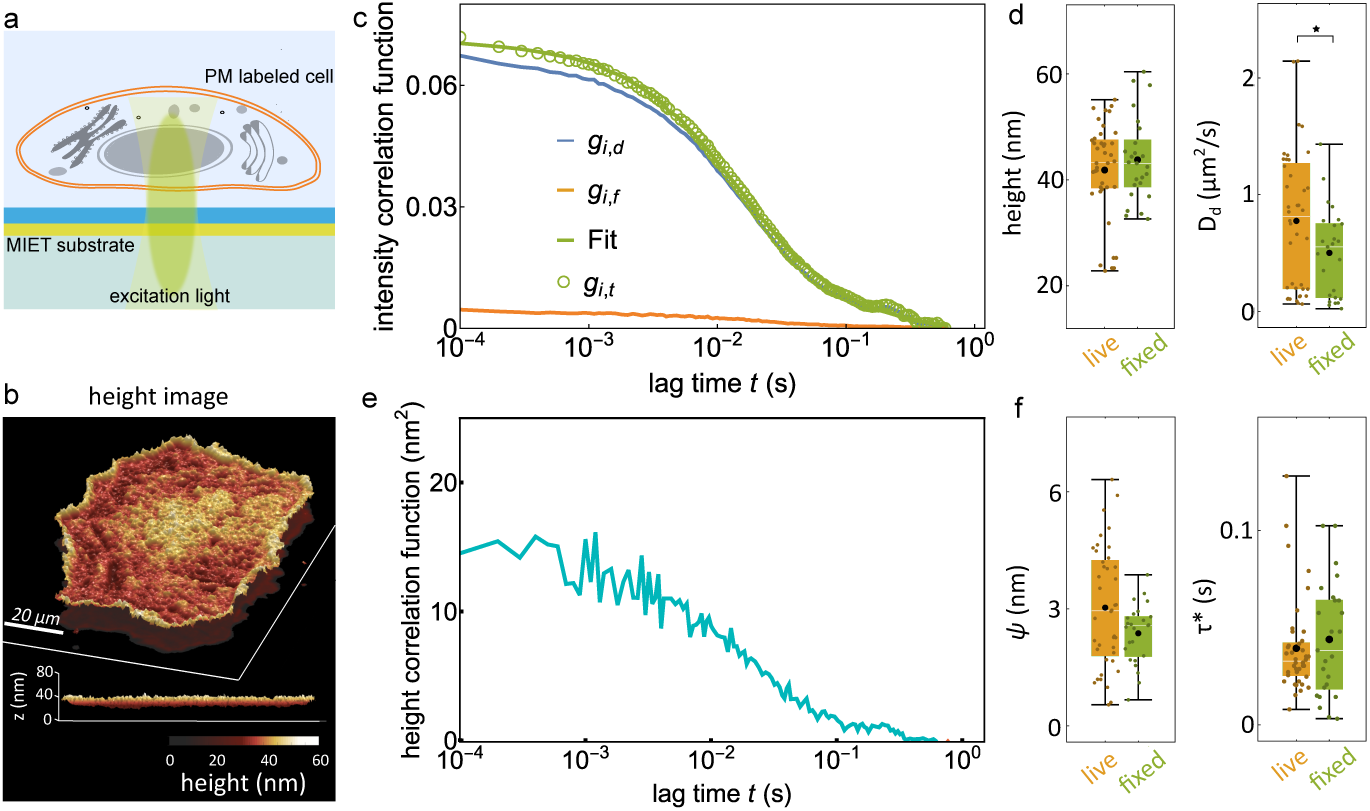
dynaMIET measurement on a cellular PM. (a) A schematic of the dynaMIET experiment conducted on a PM-labeled cell on top of a MIET substrate. (b) Reconstructed 3D topography of the fluorescently labeled basal plasma membrane of a living Cos7 cell on a MIET substrate. The bottom panel offers an alternative viewing angle. Height is color-coded for clarity. (c,e) Same as Fig. 2c,d, but for PM system. *n* = 40 independent measurements over 3 independent experiments for living cells, and *n* = 25 independent measurements over 2 independent experiments for fixed cells. (d,f) Box plots of mean height values *h*_0_, diffusion coefficients *D*, fluctuation amplitudes *ψ*, and relaxation times *τ*∗ for living and fixed cells.

Fig. 3c shows that the average height values of the membrane appear to be unaffected by the biological activity of a cell, measuring 41.9 ± 9.2 nm (mean ± SD, *n* = 40) in living cells and 43.8 ± 7.9 nm (mean ± SD, *n* = 25) in fixed cells (Fig. 3d). Notably, faster lipid diffusion was observed in living cells, with a 1.5-fold increase compared to fixed cells. Specifically, the diffusion coefficients were 0.50 ± 0.38 µm^2^/s for fixed cells and 0.77 ± 0.59 µm^2^/s for living cells (Fig. 3d). PM undulations were minimal in both living and fixed cells (Fig. 3e). Living cells exhibited only ∼1.3 times larger undulation amplitudes (3.03 ± 1.54 nm) and ∼1.4 times slower fluctuation times (28 ± 23 ms) than fixed cells (2.37 ± 0.75 nm and 39 ± 24 ms, respectively) (Fig. 3f). Considering that cross-linked proteins in fixed cells hinder active dynamics, our observations suggest that both active (energy-driven) and passive (thermal fluctuations) processes contribute to basal PM dynamics, in agreement with previous findings using alternative techniques (*57*). The increase in the fluctuation time after cell fixation implies a stiffening of the PM (*58*).

### 3D dynamics of endoplasmic reticulum

The endoplasmic reticulum (ER) is a subcellular organelle responsible for numerous essential cellular processes. Within living cells, the ER exhibits high dynamism, continuously undergoing shape remodeling. Although the relationship between ER dynamics and function is not fully understood, recent findings associate abnormal ER dynamics with various diseases (*59*). Previous reports have categorized ER dynamics in mammalian cells into three types: oscillations of established network elements, dynamics of particles within the ER lumen or membrane, and generation of new network elements (*59*). Quantifying ER dynamics remains a formidable challenge due to the complex nature of its movement.

We performed a dynaMIET measurement on ER in COS7 cells on a MIET substrate with a 10 nm silica spacer (Fig. 4a). For live-cell imaging, the ER was labeled with the commercial dye ER-Tracker Red. In Fig. 4b, we present 3D reconstructed height maps of the labeled ER in a living cell. The distance between the substrate and the ER exhibits considerable variation, ranging from 50 nm to 120 nm. It is important to note that the signal observed from a confocal microscope represents a spatial average over the size of the excitation focus (approximately 300 nm) and over vertical regions of around 500 nm (Fig. 4a). Therefore, the height images in Fig. 4b may not depict a single ER bilayer, rather an average value across the confocal volume. Notably, some locations in Fig. 4b show very low height values (approximately 50 nm), which can be attributed to contact sites between the ER and the PM (*60*). During ER point measurements, a fraction of the detected fluorescence originates from ER regions that are inside the confocal detection volume but outside the MIET-sensitive range (≥ 200 nm from the surface), contributing non–distance-sensitive background to the signal. This can significantly impact the measurement results of ER dynamics within the MIET working range. To minimize the impact of out-of-MIET ER contributions and the presence of multiple ER sheets in the focus of excitation, we excluded data that show an average ER height greater than 65 nm (see Fig. S4 and Supplementary Text).

**Figure 4:**
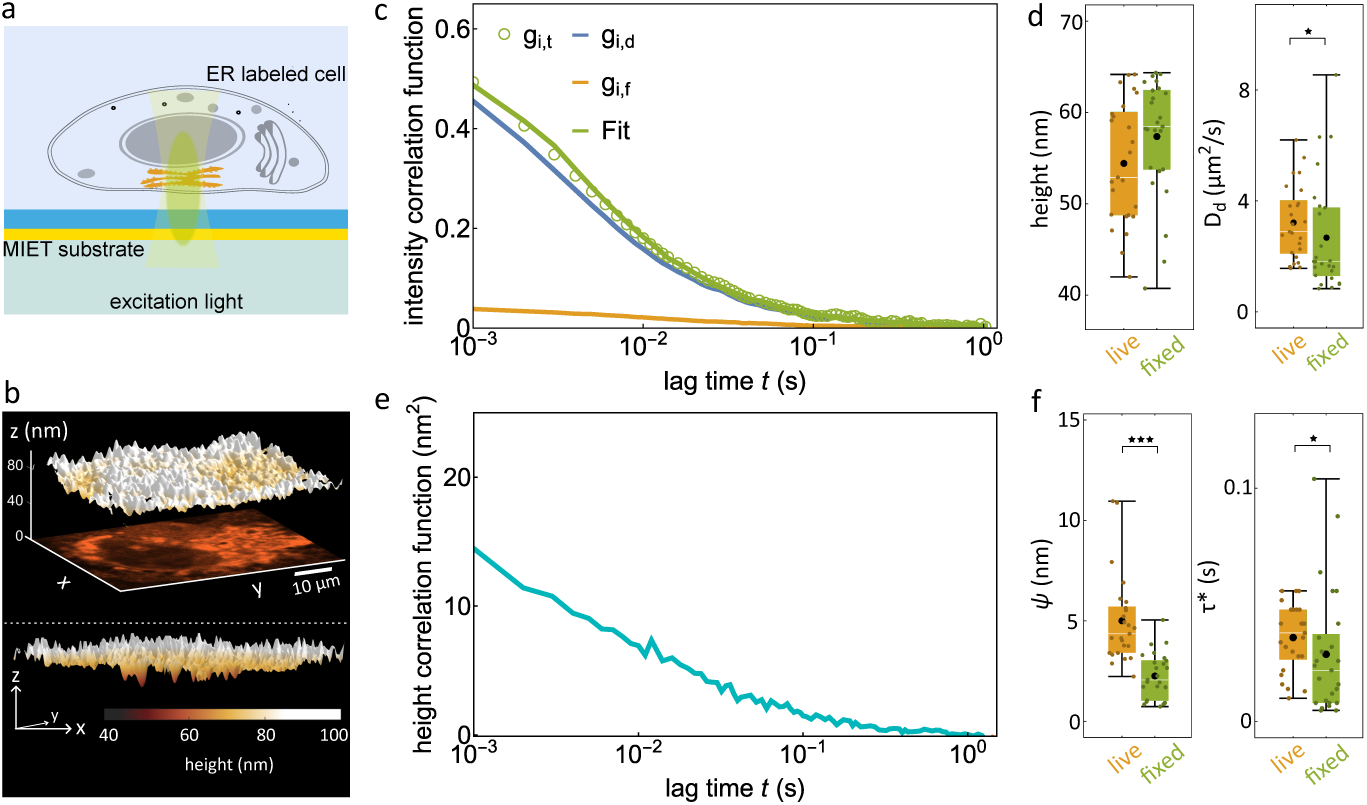
dynaMIET measurement on ER. (a,c-f) Same as Fig. 2, but for ER system. *n* = 25 independent measurements over 3 independent experiments for both living cells and fixed cells. (b) Exemplary 3D reconstructions (height maps) of fluorescently labeled ER in a live COS7 cell on a MIET substrate. The *xy*-image shows the fluorescence intensity. Bottom panel shows the height image from a different viewing angle.

The results shown in Fig. 4c-f show that we can distinguish between the lateral diffusion of lipids from the perpendicular membrane undulation components in the dynamics of the ER. Comparative dynaMIET experiments were conducted on living and fixed cells. Our findings do not reveal a significant difference between the average membrane heights *h*_0_ in both cases, with measured values of 54.4 ± 6.7 nm for living cells and 57.4 ± 6.5 nm for fixed cells (mean ± SD, *n* = 25 for living cells and *n* = 25 for fixed cells, Fig. 4d). However, significant differences are observed for all other parameters. The fluctuation amplitude *ψ* exhibits a 50% decrease in fixed cells compared to living cells, measuring 5.0 ± 2.2 nm for living cells and 2.3 ± 1.2 nm for fixed cells (Fig. 4f). The diffusion coefficient decreased by approximately 17% after fixation, from 3.2 ± 1.3 µm^2^/s in live cells and 2.7 ± 2.0 µm^2^/s in fixed cells. Finally, the correlation relaxation time *τ*^∗^ showed a 20% decrease from 36 ± 14 ms in living cells to 28 ± 26 ms in fixed cells (Fig. 4f). The substantial reduction in the amplitude of undulation *ψ* from living cells to fixed cells can be attributed to the fact that the dynamics of ER is predominantly determined by the active motion of the cytoskeleton network and motor proteins (*61, 62*).

### 3D dynamics of the nuclear envelope

The nuclear envelope (NE) serves as the physical boundary between the cytoplasm and the nucleoplasm, separating two highly dynamic systems: the cellular cytoskeleton and nuclear chromatin. We performed dynaMIET measurements on live and fixed COS7 cells expressing eGFP fused to the nuclear pore protein Nup153, see Fig. 5a and Fig. S5. Fig. 5b shows the measured height profile of the proximal part of the nuclear envelope, showing an average height value of approximately 130 nm well within the working distance of MIET.

**Figure 5:**
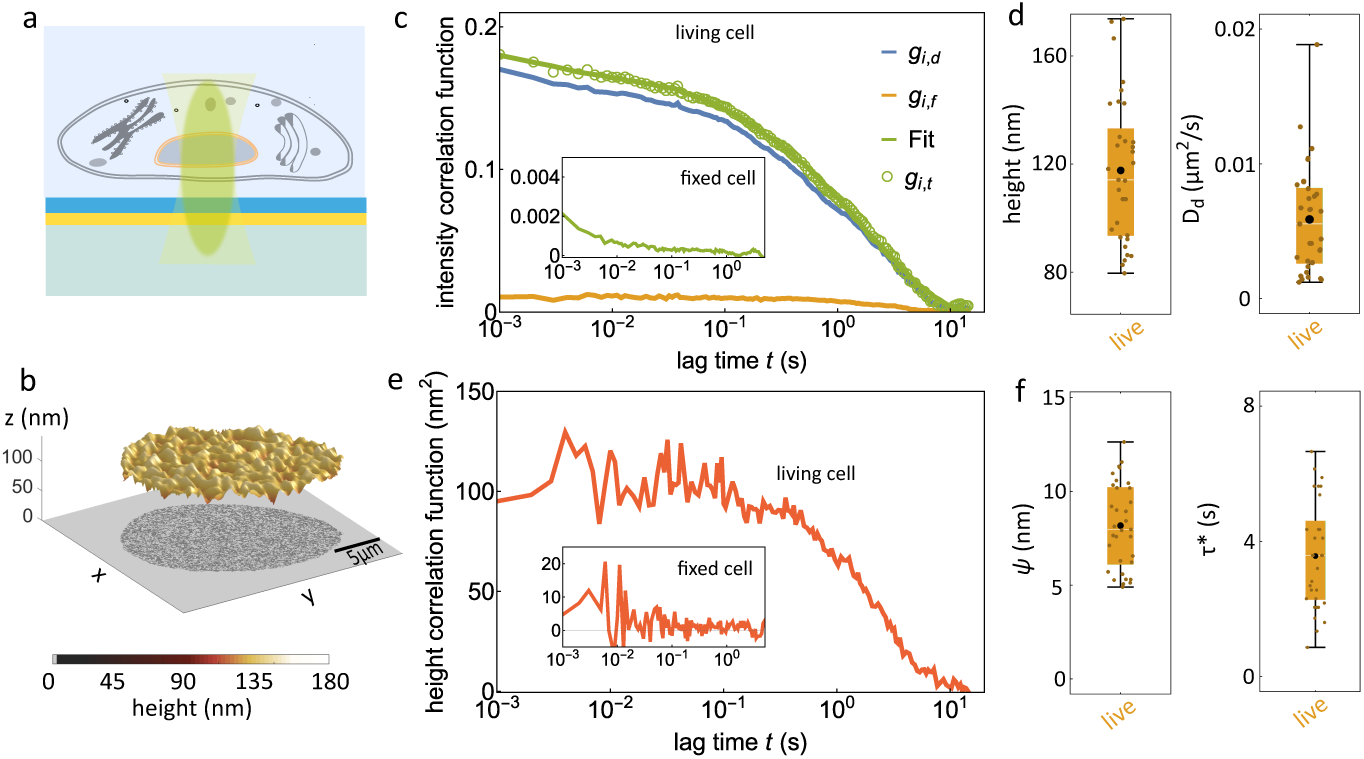
dynaMIET measurement of NE dynamics. (a-f) Same as Fig. 3, but for NE system. (c) iACFs obtained for living eGFP-nup153-modified COS7 cell. The green solid line represents a fit with eq S3. Inset shows measured *g*_*i*_ for a fixed cell. (e) Calculated hACF for a living and a fixed cell (inset).

Fig. 5c shows iACFs for NE Nup153 lateral diffusion and membrane undulations in living cells. The observed diffusion is ca. 100 times slower than values measured for lipid diffusion in deflated GUVs, with a diffusion coefficient of only 0.0059 ± 0.0040 µm^2^/s (*n* = 33, Fig. 5d). The relaxation time of NE membrane undulations is also 60 times slower than that for deflated GUV, determined to be on the timescale of seconds (3.6 ± 1.6 s, Fig. 5f), in good agreement with previous measurements (*63, 64*). The slower dynamics of the NE compared to other cellular membranes can be attributed to its double-bilayer structure, and to strong links to the cytoskeleton on the cytoplasmic side and to the chromatin on the nucleoplasmic side.

Our results show no diffusion or undulations of NE in fixed cells (Fig. 5c,e inset), where the amplitude of the measured total iACF is only 0.002 or 100 times smaller than its value measured in living cells. This is attributed to the heavy cross-linking between the NE and the cytoskeleton and nuclear proteins in the fixed cells. The small residual iACF amplitude may be due to triplet-state photophysics of the eGFP labels. The hACF is too noisy to allow for determining reliable values of relaxation time and amplitude.

## Discussion

We introduced dynaMIET as a robust and accessible method for probing the 3D dynamics of biological membranes in both *in vitro* and *in cell* systems. By integrating the nanometer axial precision of MIET with the microsecond temporal resolution of FCS, dynaMIET enables simultaneous quantification of lateral diffusion and vertical membrane undulations. This approach is straightforward to implement on any standard confocal microscope with FLIM modality, and crucially, it requires no empirical calibration, as the necessary calibration curves can be directly accurately computed from theoretical models.

We demonstrated the versatility of dynaMIET through a wide range of applications, including measurements on GUV membranes, and several cellular membrane systems, such as PM, ER, and NE. From these measurements, we extracted a suite of key biophysical parameters, including diffusion coefficients, fluctuation amplitudes, relaxation times, membrane tension, and membrane–substrate interaction strengths. These insights are particularly valuable for understanding how the mechanical environment, such as substrate adhesion or cytoskeletal coupling, influences membrane organization and function. Beyond model membranes, we demonstrated the usefulness of dynaMIET for membrane studies in living and fixed cells.

dynaMIET offers significant scalability and flexibility. It can transition from a point measurement setup to fast-scanning FCS or camera-based wide-field fluorescence lifetime imaging (*65*). This would enable spatially resolved mapping of membrane dynamics across large cellular areas. Furthermore, combining dynaMIET with dual-labeling strategies, where both membrane lipids and proteins are independently labeled, opens exciting possibilities for probing protein–membrane interactions at nanometer scales. This includes tracking how peripheral or transmembrane proteins modulate local membrane curvature, tension, or lipid mobility, which are critical aspects in signaling, vesicle trafficking, and organelle biogenesis. We anticipate that dynaMIET will evolve into a potent and versatile tool for investigating membrane biophysics, particularly in the realms of nanoscale structure and microsecond fluctuations. This methodology holds promise for diverse applications and contexts within the field.

## Acknowledgments

We thank Anna Chizhik, Ingo Gregor, and Alexey Chizhik for their help and support.

## Author contributions

J.E. and T.C. conceived the project. T.C. prepared all samples of GUVs and SLBs. J.I.G. prepared most of the samples of cells and all genetically modified cells. T.C. and J.I.G. performed all the measurements. T.C. analyzed all the data and made all figures both in the main text and supplementary information. D.W prepared some cell samples for PM measurement. N.K. wrote part of MATLAB code to analyze the data and simulation. T.C. and J.I.G. helped with preparing the manuscript, which was written by J.E.

## Data and materials availability

All code used for generating Fig. s 1-6 are deposited on GitHub at https://gitlab.gwdg.de/tchen1/3dmembranedynamics.git. In particular, the depository contains:

- *Matlab* (v. 2022b, MathWorks^®^ Inc.) code used for calculating the MIET and brightness calibration curves, intensity trace, height trace, intensity autocorrelation function, height autocorrelation function, as well as fitting the intensity autocorrelation function, height autocorrelation function;
- *Mathematica* (v. 13.2.1.0 Wolfram Research Inc.) notebook that generates the graphs of all figures;
- A Readme file to explain all analysis detials.

The 3D height profiles were calculated using a freely available MATLAB-based MIET-GUI software (*66*).

## Supplementary materials

Materials and Methods

Supplementary Text

Figs. S1 to S6

