## Supplemental figures and text for "Quantifying 3D Live-Cell Membrane Dynamics Using Dynamic Metal-Induced Energy Transfer Spectroscopy (dynaMIET)"

**This PDF file includes:**

Materials and Methods

Supplementary Text

Figures S1 to S7

### Materials and Methods

#### MIET theory

This fluorophore-metal interaction can be modeled in a semi-classical approach based on Maxwell's equations. For this purpose, one considers the fluorescent molecule as an ideal electric dipole emitter and calculates its electromagnetic field in the presence of the MIET (GIET) substrate, as detailed in Refs. (*67, 68*). This allows for the calculation of the total dipole emission rate (inverse radiative decay rate of the excited fluorophore) as a function of dipole position, orientation (expressed by the angle $\beta$ between its dipole axis and the vertical direction) and geometry and electrodynamic properties (complex-valued refractive indices) of the substrate. Together with the knowledge of the fluorescence quantum yield $\phi$ (the ratio of radiative to total transition rate), one then obtains values of the observable fluorescence lifetime $\tau_f(h,\beta)$ and brightness $b(h,\beta)$ of a fluorophore as a function of its distance $h$ and orientation angle $\beta$ as

$$\frac{\tau_f(h,\beta)}{\tau_0} = \frac{S_0}{\phi S(h,\beta) + (1-\phi)S_0} \tag{S1}$$

$$b(h,\beta) \propto \int_0^{\pi/2} d\beta \sin\beta \frac{N(h,\beta)}{S(h,\beta)} C(h,\beta) \tag{S2}$$

In Eq. S1, $S(h,\beta)$ represents the total emission power of the emitter, where $S_0$ stands for the emission power of an electric dipole emitter with unity quantum yield in a homogeneous unbounded dielectric medium with a refractive index $n$, where the emitter has fluorescence lifetime $\tau_0$ (free-space fluorescence lifetime). Moving to Eq S2, $N(h,\beta)$ is the total far-field emission radiated into the lower half space (glass coverslip), while $C(h,\beta)$ signifies the relative proportion of emission collected by the objective lens. The angular dependence of $S(h,\beta)$ can be decomposed as $S(h,\beta) = S_\perp(h)\cos^2\beta + S_\parallel(h)\sin^2\beta$, where $S_\perp(h)$ and $S_\parallel(h)$ are the radiative emission rates of emitters oriented perpendicular and parallel to the substrate, respectively. This decomposition extends similarly to $N(h,\beta)$ and $C(h,\beta)$. A detailed mathematical explanation is given in Ref. (*69*). It is crucial to note that the functions $S_\perp(h)$ and $S_\parallel(h)$ are solely dependent on $h$.

#### Simulation of membrane diffusion and fluctuation near a gold surface.

To investigate the fluorescence signal modulated by membrane dynamics near a plasmonic interface, we performed numerical simulations of fluorescent particle diffusion on a fluctuating lipid membrane positioned above a gold surface. The lateral diffusion of particles was modeled as two-dimensional Brownian motion within a $50\,\mu$m $\times$ $50\,\mu$m box with periodic boundary conditions, using a diffusion coefficient $D = 5.0\,\mu\text{m}^2/\text{s}$, particle surface concentration $C = 40\,\text{particles}/\mu\text{m}^2$, and a Gaussian excitation beam with a waist of $\omega = 0.3\,\mu$m. Membrane height fluctuations were simulated using an Euler Mayurama method with a mean height $z_0 = 40\,$nm, standard deviation $\sigma_z = 10\,$nm, and correlation time $\tau_c = 50\,$ms, reflecting thermal motion within a constrained potential. Each simulation included $10^5$ time steps with a temporal resolution of $\Delta t = 0.1\,$ms, for a total simulated duration of 10 seconds per run. The process was repeated 10 times to obtain averaged statistical results. Fluorescence intensity at each time point was computed by summing the Gaussian-weighted contributions of all particles within the focal area, modulated by the instantaneous membrane height using precomputed MIET brightness and lifetime calibration curves. The MIET parameters were: free-space fluorescence lifetime $\tau_{\text{free}} = 1\,$ns, fluorophore quantum yield $QY = 0.8$, emission wavelength $\lambda = 700\,$nm, and a multilayer structure composed of 2 nm Ti / 10 nm Au / 1 nm Ti / 10 nm $SiO_2$. Autocorrelation functions were calculated from the resulting time traces to disentangle the contributions of diffusion and vertical membrane fluctuations to the observed FCS signal.

**dynaMIET analysis flow chart**

Measured fluorescence intensity fluctuations can be analyzed by calculating an intensity autocorrelation function $g_{i,t}(t)$ (iACF). The value of the iACF for a lag time $t$ is defined as the time average of the product of intensity deviations from the mean value at time, $t'$ $\delta I(t') = I(t') - \langle I \rangle$, and at time $t' + t$, $\delta I(t' + t) = I(t' + t) - \langle I \rangle$, i.e. $g_i(t) = \langle \delta I(t')\delta I(t' + t) \rangle$, where the angular brackets denote averaging over time $t'$. As mentioned in main text, in the presence of a MIET substrate, the total iACF $g_{i,t}(t)$ can be divided into a part $g_{i,d}(t)$ related to the lateral diffusion of single lipids, a part $g_{i,f}(t)$ related to vertical undulations of the membrane, and a part $g_{i,tri}(t)$ related to the triple state relaxation of the fluorophore (see eq. S1).

We separated the $g_{i,d}(t)$ and $g_{i,f}(t)$ by so-called time-tagged time-resolved (TTTR) single-photon counting. For each detected photon, TTTR provides two timestamps: the macrotime that counts the number of laser pulses between the start of the measurement and the photon detection, and the microtime that is the delay time between the last laser pulse and the photon detection. The macrotime is utilized for computing intensity traces and the iACF $g_{i,t}$, while the microtime is used to calculate the fluorescence lifetime.

In our dynaMIET analysis (Fig. S7), we start with computing an intensity trace $I(t)$ over time bins with width Δ t = 100 μs or 1ms, which is then used to calculate the iACF $g_{i,t}(t)$ (Fig. S7i, after photobleaching correction, see Fig. S1 and Supplementary Text). Subsequently, we determine the fluorescence lifetime within each time bin by computing the square root of the variance of the microtimes recorded during each time bin (Fig. S7ii). An ideal fast lifetime estimation should have less than 10 % uncertainty, which requires approximately 100 photons (*70, 71*). This determines the minimum width of the time binning: one needs approximately 100 photons per time bin for a reasonable lifetime estimate. Taking into account that the maximum photon detection rate that a single photon detector can handle is typically around 4,000 kcps (kilo counts per second), one obtains a minimal possible time bin width of 25 μs. However, the photon detection rate varies as a function of fluorophore brightness and concentration, which results in a varying temporal resolution for different samples (see Supplementary Text and Fig. S2).

In a next step, each fluorescence lifetime per time bin is converted into a height value using the lifetime-versus-height calibration curve (Fig. 1c, blue curve), resulting in a continuous trace of height values $h(t)$ (Fig. S7iii). Furthermore, relying on the brightness-height calibration curve (Fig. 1c, yellow curve), a relative brightness trace ($br(t)$) is obtained (Fig. S7iv), facilitating the direct computation of the MIET-induced height-related correlation function ($g_{i,f}(t) = \langle br(t')br(t'+t)\rangle_{t'} / \langle br(t)\rangle^2$ ). For $g_{i,d}(t)$, the diffusion can be described with a 2D diffusion model, $g_{i,d} = 1/(N(1+t/t_D))$, where N is the total molecules in the confocal volume and $\tau_D$ is the diffusion time of lipids through the confocal volume Therefore, from Eq. 1, combining all the contributions, the iACF $g_{i,t}(t)$ can be fitted with following model:

$$g_{i,t}(t) = \left[1 + \frac{F\exp(-t/\tau_T)}{1-F}\right] \frac{g_{i,f}(t)}{N(1+t/\tau_D)} \tag{S3}$$

where $F$ and $\tau_T$ are the triplet state amplitude and time that take into account the triplet state pumping and decay of the used fluorophores. Finally, the diffusion coefficient $D$ is determined from $D = w^2/(4\tau_D)$, where $w$ is the beam waist of the laser excitation focus in the plane of the membrane.

The height trace $h(t)$ provides information about the vertical undulations of a membrane.

It can be decomposed into a mean value $h_0$ plus a deviation from this mean value $\delta h(t)$, i.e. $h(t) = h_0 + \delta h(t)$. The temporal dynamics of these membrane undulations can be characterized by the height autocorrelation function $g_h(t) = \langle \delta h(t') \delta h(t' + t) \rangle_{t'}$ (hACF).

Next, we can utilize the hACF $g_h(t)$ and the diffusion-related iACF $g_{\mathrm{dif}}(t)$ to analyze the 3D dynamics of the membrane. The amplitude of the hACF $g_h(t)$ determines the mean displacement amplitude ($\psi = \sqrt{< \delta h^2 >}$), and the decay time of the hACF, determined by the point where this function has decreased to half its maximum value, yields a relaxation time $\tau^*$.

**dynaMIET measurement**

dynaMIET measurements were conducted using a custom-built confocal microscopy setup. The excitation sources comprised a pulsed diode laser ($\lambda_{exc}$ = 640nm, LDH-D-C 640, PicoQuant) for red dye and a white laser (NKT Photonics, Koheras SuperK Power) with Acousto-Optic Tunable Filters (AOTFnC-400.650-TN, Pegasus Optik Gmbh). The beam was collimated from an infinity-corrected 4x objective (UPISApo 4X, Olympus), then reflected by a dichroic mirror (Di01-R405/488/561/635, Semrock) towards a high numerical aperture objective (UApoN 100X, oil, 1.49 N.A., Olympus). Emission light was focused into a pinhole with a diameter of 100 µm and refocused onto two avalanche photodiodes ($\tau$-SPAD, PicoQuant). A long-pass filter (BLP01-647R-25, or LF561/LP-D-000, or LF488/LP-C-000, Semrock) and two band-pass filters (Brightline HC692/40, or FF01-609/54-25, or FF01-525/45-25, Semrock) were positioned before the pinhole and the detector, respectively. Single-count data from the detector were processed using a multi-channel picosecond event timer (Hydraharp 400, PicoQuant). Imaging scanning was facilitated by a fast Galvo scanner (FLIMbee, Picoquant).

For live cell measurements, a microscopy-based incubator (ibidi Stage Top Incubation System-Blue Line, ibidi) maintained controlled conditions (temperature: 37 °C, $CO_2$ concentration: 5%, humidity: 40%). In a typical measurement, the sample was initially scanned at the MIET substrate surface to determine the contour of the GUVs or cells. Subsequently, an appropriate point position was selected for single-point acquisition, and data were recorded for at least 2 minutes for each data point. The sample within the working distance of MIET can be determined by its fluorescence curve (Fig. S6).

### Substrate preparation

We followed the procedures outlined in our previous publications (*46, 68*) to prepare the gold-modified substrate. A layer-by-layer deposition technique was used, where 2 nm of titanium, 10 nm of gold (or 40 nm for SLB measurement), 1 nm of titanium, and 10 nm of $SiO_2$ were successively electron-beam evaporated onto the surface of a glass coverslip. The electron-beam evaporation was carried out using a Univex 350 electron beam source (Leybold) under high-vacuum conditions (approximately $10^{-6}$ mbar). To ensure maximal homogeneity, the deposition was conducted at the slowest rate (1 $\mathring{A}s^{-1}$). The spacer thickness was continuously monitored during the evaporation process using an oscillating quartz unit. This gold-covered substrate is referred to as the MIET substrate.

Prior to the MIET experiment, the MIET substrate underwent plasma cleaning for 60 seconds at the highest plasma intensity (Harrick Plasma). Subsequently, it was incubated in a 5 mg/mL$^{-1}$ solution of bovine serum albumin (BSA, Sigma-Aldrich) for 15 minutes to prevent nonspecific interactions in GUV experiments.

### Vesicles and SLB preparation

SLB was prepared through the fusion of small unilamellar vesicles (SUVs), which were generated using an extrusion method. A 100 μL mixture containing 10 mg/mL 1,2-Dioleoyl-sn-glycero-3-phosphocholine (DOPC) lipids and 0.05 mol% DPPE-atto655 lipids in chloroform was dried in a vacuum for 1 h at 30 °C. The resulting lipid film was then re-dispersed in 500 μL of PBS buffer (pH = 7.4) and shaken for 1 h at 37 °C. SUVs were obtained by extruding the solution through a polycarbonate filter (Whatman) with a 50 nm pore diameter for 15 cycles. The SLB was prepared by incubating the SUV solution on the MIET substrate for 1 h at room temperature, followed by washing with PBS buffer.

GUVs made of SOPC were fabricated using the electroformation method (*68*). A 100 μL lipid mixture containing 10 mg/mL SOPC, 2 mol% 1,2-dioleoyl-sn-glycero-3-phosphoethanolamine-N-(methoxy(polyethylene glycol)-2000) (DOPE-PEG2000), and 0.05 mol% DPPE-atto655 was deposited on two steel electrodes and evaporated for 3 h under vacuum at 30 °C. The electrodes were sealed with a Teflon chamber and filled with 500 μL of sucrose solution (230 mM). The chamber was subjected to an alternating electric current at 15 Hz for 3 h with a peak-to-peak voltage of 1.6, followed by 8 Hz for 30 min. After electro-formation, the GUVs suspension was stored at −4°C and used within 3 days. The GUVs suspension was diluted 50 times in PBS buffer (400 mOsml$^{-1}$, 10mM $Na_2HPO_4$, 2mM $KH_2PO_4$, 3 mM KCl, and 180 mM NaCl, pH 7.4) for the observation of floppy GUVs. For all GUV experiments, the diluted GUVs solution with observation buffer was added to a chamber (Culture-Insert 4 Well, ibidi GmbH) on the MIET substrate. A glass slide covered the chamber, and the setup was left undisturbed for 10 min at room temperature for equilibration before the measurement.

#### Cell culture and mambrane labeling

COS7 monkey kidney cells were cultured at 37°C in 5% CO2 in T-75 culture flasks using Dulbecco's Modified Eagle's Medium (DMEM) - high glucose (Sigma, D6429-500L) supplemented with 10% fetal bovine serum (Sigma-Aldrich, F7524) and 1% penicillin/streptomycin (Sigma, P4333). Cells were seeded onto 35-mm glass-bottom dishes (Ibidi, 81158), either gold-coated or uncoated, at 8 × $10^4$ cells per dish and cultured for 24 h. For experiments using fixed cells, samples were washed with PBS and fixed in 4% paraformaldehyde/PBS solution for 20 minutes.

ER-Tracker Red (0.5 μM solution in DMEM) and CellMask Deep Red (500 ng/ml solution in DMEM) were used for specific labelling of the endoplasmic reticulum (ER) and plasma membrane (PM), respectively. For nuclear envelope labeling, cells were transfected one day before imaging with the pEGFP(C3)-Nup153 plasmid encoding fluorescent nucleoporin-153 (*72*) using Effectene Transfection Reagent (Qiagen, 301425). pEGFP(C3)-Nup153 was a gift from Birthe Fahrenkrog (Addgene plasmid # 64268; http://n2t.net/addgene:64268; RRID:Addgene_64268).

### Supplementary Text

#### Supporting Text 1. Correction of the photobleaching

The photobleaching of fluorophores during the measurement significantly impacts the fluorescence intensity autocorrelation function gi,t (see Fig. S1). In our experiments, we noticed the photobleaching effects in the fluorophores of CellMask Deep Red and eGFP protein. As shown in Fig. S1, a discernible decay component is evident in the correlation curve, attributed to photobleaching.

To mitigate the influence of photobleaching on gi,t, we performed a correction on the intensity time trace (I(t)) by applying a bi-exponential decay function:

$$z_{fit} = \sum_{i=1}^{2} a_i e^{-I(t)/t_i} + b \tag{S4}$$

where $z_{fit}$ denotes the intensity decay curve, $a_i$ represents the amplitudes, $t_i$ refers to the decay constants, and b signifies an intensity offect. Consequently, we derived the relative intensity trace $I_{rel}(t) = I(t)/z_{fit}$ (Fig. S1). As illustrated in Fig. S1, the decay component attributed to photobleaching is effectively eliminated post-photobleaching correction.

**Supporting Text 2. Determination of temporal resolution**

The temporal resolution of measurement is contingent upon the count rates, which directly influence the accuracy of the lifetime and subsequent height calculations According to previous studies, reliable accuracy in fast lifetime estimation necessitates a minimum of approximately 100 photon counts (*70, 71*). The photon count in each bin of the intensity trace is determined by factors such as fluorophore concentration, brightness, and excitation intensity. In our experiments involving four samples (GUV, PM, ER, NE), the count rates can easily exceed 1,000 kilo counts per second (k.c.p.s.) for GUV and PM samples. However, for ER and NE measurements, the count rates are lower, approximately 500 k.c.p.s. and 200 k.c.p.s., respectively. To ensure the accuracy of lifetime calculation, we employed a time bin width of 100 $\mu$s for GUV and PM samples, corresponding to approximately 100 photon counts per bin. For ER and NE measurements, we utilized a time bin width of 1 ms, corresponding to approximately 500 and 200 photon counts per bin, respectively. To assess the effect of photon count on the accuracy of lifetime and height calculations, we generated intensity traces (see Fig. S2) and their corresponding height traces (Fig. S2) using different time bins ranging from 25 $\mu$s to 1 ms. As demonstrated in Fig. S2c and d, the intensity autocorrelation functions (iACFs) remained unaffected by the bin width. However, the height autocorrelation function (hACF) showed dependency on the bin width. Specifically, the amplitude of the hACFs did not vary significantly when the photon counts per bin exceeded 40. Conversely, when the photon count dropped to 20 or lower, the amplitude of the hACF decreased substantially compared to other scenarios, resulting in an error of more than 1 nm in the membrane fluctuation amplitude.

**Supporting Text 3. Statistics of ER dataset**

When conducting a point measurement with a confocal microscope on the ER, positions are randomly selected to collect data. However, due to the sac-like and heterogeneous structures of ER, the measured result represents a spatial average over all ER within the size of the excitation focus. Consequently, some ER regions outside the working distance of MIET are inadvertently included. This inclusion can lead to smaller measured values of fluctuation amplitude for ER, as movements form ER regions beyond the working distance of MIET do not contribute to height-related intensity fluctuations. To mitigate this effect, we implemented a lifetime threshold to exclude data points with large lifetimes, where quenching is not evident. Because lots of the ERs in these data points were placing out of the working distance of MIET, their lifetimes were dominated by these out-working-distance ER. We established this threshold based on the MIET calibration curve and brightness calibration curve of the of ER tracker Red fluorophore. As shown in Fig. S4, both calibration curves present steepest slopes for the height region below 65 nm (equivalent to 3.67 ns in lifetime), indicating the optimal working region of ER tracker Red is below 65nm. Consequently, we excluded data points with a lifetime exceeding 3.67 ns for the ER statistics. It is noteworthy that approximately 40% of data points were excluded from the statistics using this threshold.

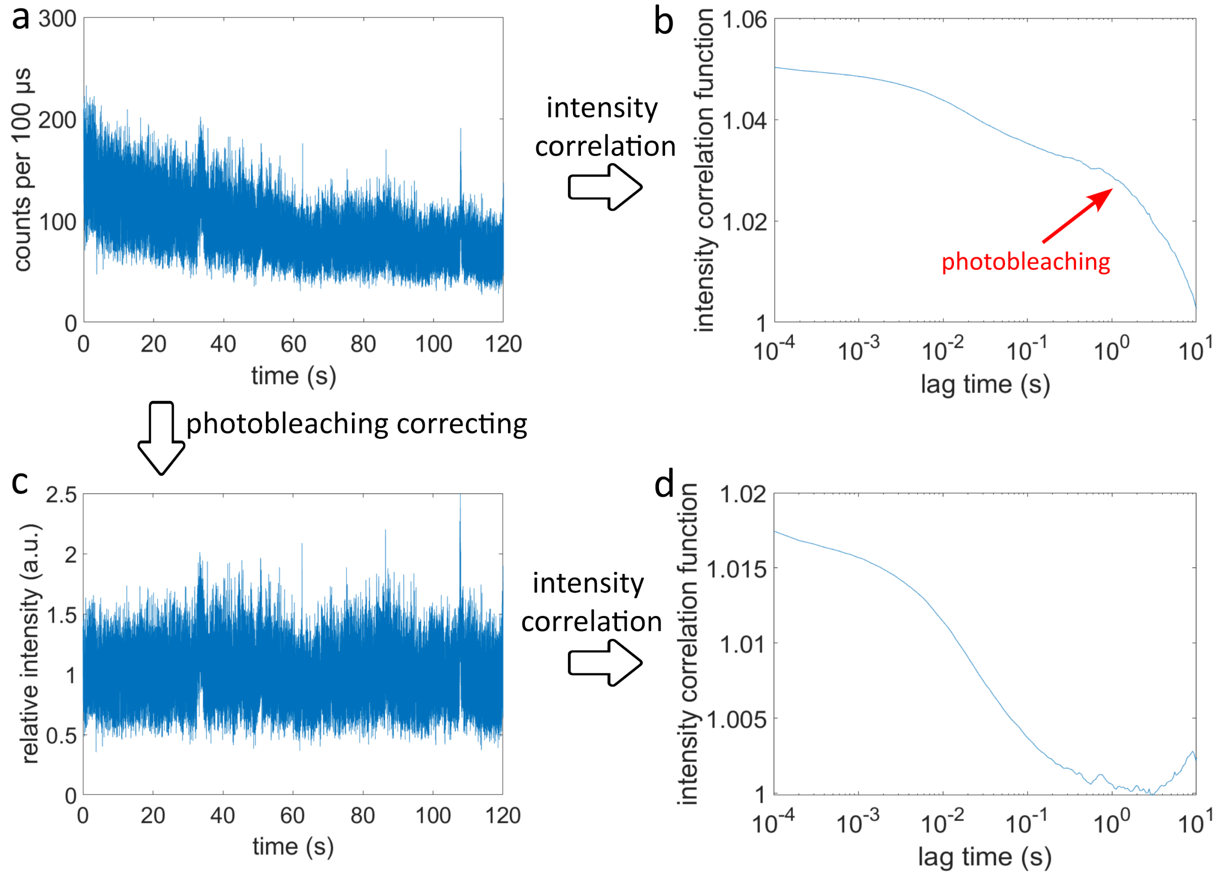


**Figure S1**: **Photobleaching correction process.** (a) Fluorescence intensity trace measured from the CellMask Deep Red labelled cell on a MIET substrate. (b) The associated intensity correlation curve is depicted without photobleaching correction. The red arrow indicates the decay component attributed to photobleaching. (c) Fluorescence relative intensity trace after the photobleaching correction. (d) The resulting intensity correlation curve is calculated from the relative intensity trace.

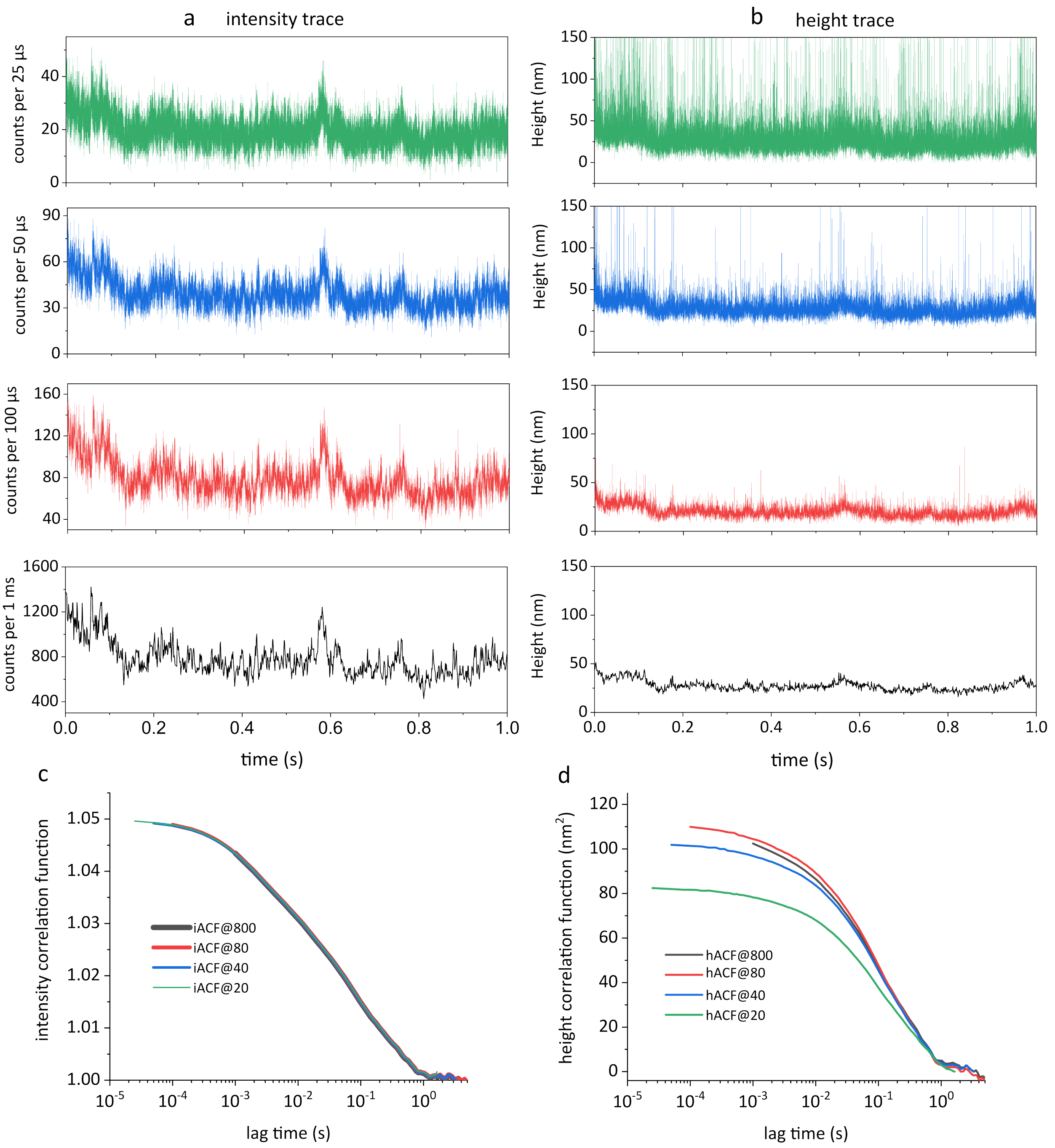


**Figure S2**: **Determination of temporal resolution.** (a) Fluorescence intensity traces calculated at different time bin width, and (b) the corresponding height traces. (c) Intensity autocorrelation functions (iACFs) are constructed from different intensity traces as depicted in panel a. The notation 'iACF@800' indicates averaging 800 photon/bin to calculate the iACF, with similar interpretations for other cases. (d) Height autocorrelation functions (hACFs) are constructed from different height traces illustrated in panel b. The amplitude of hACFs at $10^{-3}$ lag time are: 102.4 nm$^2$ (hACF@800), 104.5 nm$^2$ (hACF@80), 96.8 nm$^2$ (hACF@40), and 78.2 nm$^2$ (hACF@20), corresponding to the membrane fluctuations $\psi$ are: 10.1 nm$^2$ (hACF@800), 10.2 nm$^2$ (hACF@80), 9.8 nm$^2$ (hACF@40), and 8.8 nm$^2$ (hACF@20).

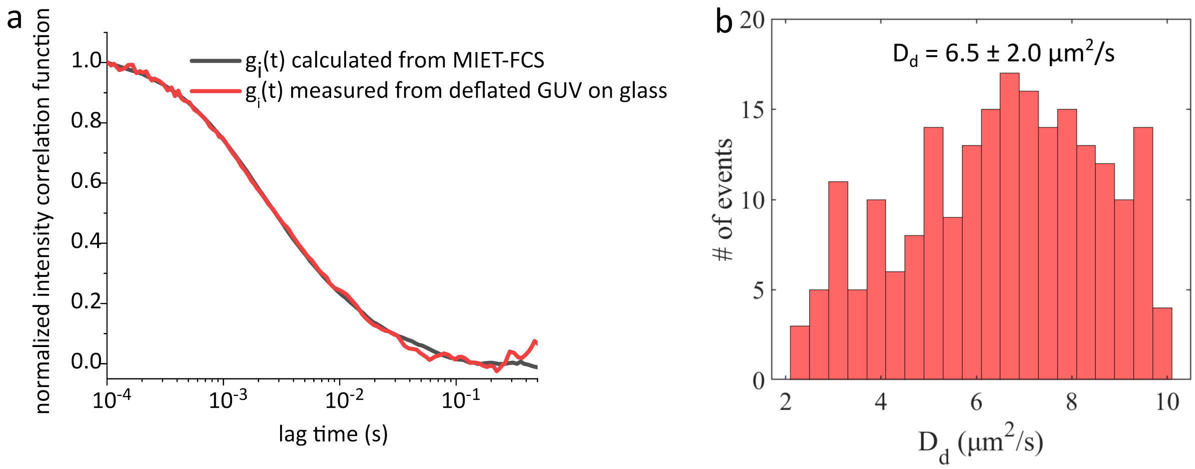


**Figure S3**: **Conventional FCS on deflated GUVs.** (a) Normalized intensity correlation function gi(t) calculated from dynaMIET, the data was measured from deflated GUV on MIET substrate (black curve), and the normalized intensity correlation function gi(t) directly measured from the deflated GUV on glass (red curve). (b) Histogram illustrating the diffusion coefficients Dd measured from the deflated GUV on glass. The mean Dd is 6.5 ± 2.0 $\mu m^2/s$ (mean ± SD, n = 214).

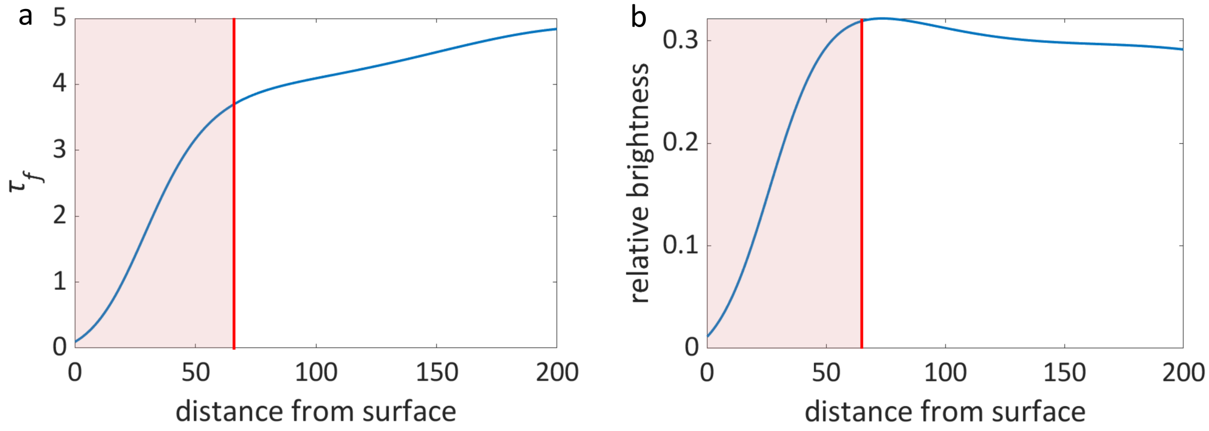


**Figure S4**: **Calculated calibration curves of dye ER Tracker Red.** (a) Calculated fluorescence lifetime of an ER Tracker Red as a function of its distance from the surface of a MIET substrate, (b) and calculated relative brightness of an ER Tracker Red as a function of its distance from the surface of a MIET substrate. The MIET substrate consists multiple layers: 10 nm silica, 1 nm titanium, 10 nm gold film, 2 nm titanium, and glass coverslip. All calculations were performed for an emitter with maximum emission wavelength of 590 nm, fluorescence quantum yield of 0.9, free-space lifetime of 4.6, and random orientation. The red lines indicate the threshold for the statistics of ER dataset. The free-space lifetime of 4.6 was measured from ER Tracker Red dye in cell on a glass surface and the quantum yield of 0.9 take the values from published reference (*73*).

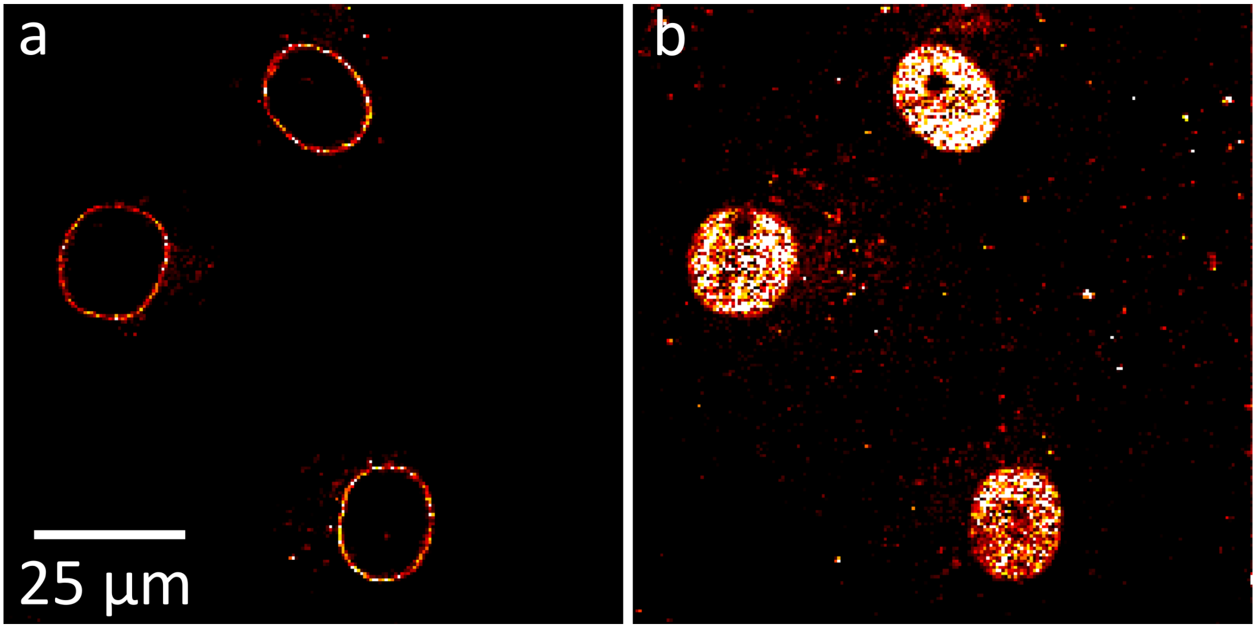


**Figure S5**: **Fluorescence image of GFP-nup153-modified cells.** (a) The image was scanned at the center of the cell nucleus. (b) The image was scanned at the bottom of the cell nucleus.

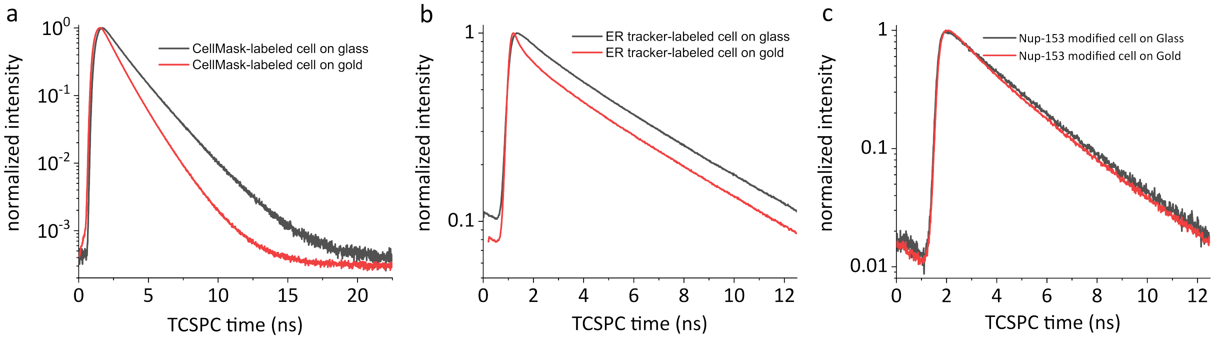


**Figure S6**: **Fluorescence lifetime decay curve for different samples.** (a) Fluorescence lifetime decay curve for CellMask-Deep-Red labelled cell on glass and on gold surface. (b) Fluorescence lifetime decay curve for ER-Tracker-Red-labelled cell on glass and on gold surface. (c) Fluorescence lifetime decay curve for eGFP-nup153-modified cell on glass and on gold surface.

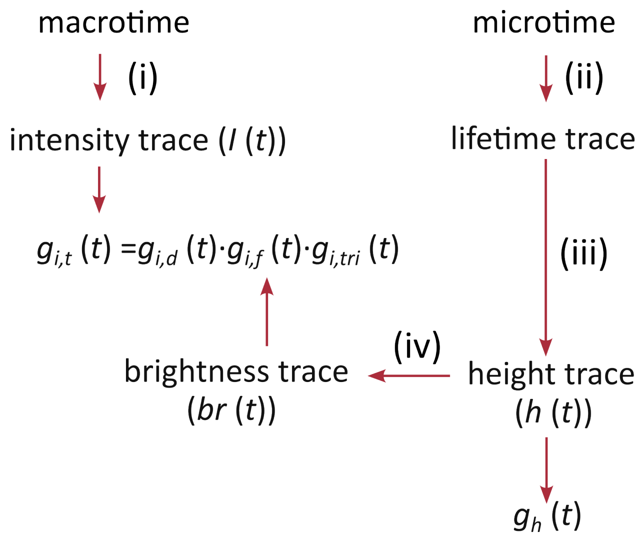


**Figure S7**: dynaMIET analysis flow chart.
